# The expression of cannabinoid-related genes in multiple disease cell lines

**DOI:** 10.1101/759332

**Authors:** Philip A. Arlen, Xingzhu Wu

## Abstract

The endocannabinoid system comprises a series of ligands and receptors that have putative roles in regulating physiological and cognitive processes such as pre- and post-natal development, appetite, pain sensation, mood, and memory. Besides the endocannabinoids, these endogenous receptors are also capable of mediating the pharmacological effects of phytocannabinoids, which are derived from the cannabis plant. We sought to interrogate a panel of cell lines, representative of multiple human diseases, to characterize their cannabinoid-mediating gene expression. We found all lines expressed one or more gene product that rendered the cell potentially responsive to cannabinoids. Moreover, the expression profiles differed between normal and cancerous cells, as well as between cells derived from the brain, pancreas, and skin. Taken together, given the presence of one or more of these mediating gene products, our findings suggest a potential therapeutic role for phytocannabinoids.

## Introduction

The cannabis plant (*Cannabis sativa* L.) contains a large number of pharmacologically active compounds called phytocannabinoids. The most abundant of these compounds are Δ9-tetrahydrocannabinol (THC) and cannabidiol (CBD), although the amounts and proportions of the various phytocannabinoids in each plant vary by strain and can be adjusted by breeding.

Many cannabinoids, including CBD, have been shown to be non-toxic and to possess anti-tumor activity in multiple cancer types ^1^. Numerous mechanisms have been proposed to explain these anti-tumor effects, including the production of reactive oxygen species ^2,3^, promotion of apoptosis ^4^, a reduction in inflammation ^5^, activation of receptors that may inhibit drug transport ^6^ and tumorigenesis ^7^, and potentiating the activity of multiple types of chemotherapy ^8^.

In spite of its wide spectrum of pharmacological activities, little is known about the molecular mechanism of cannabinoid action. Several gene products, however, have been implicated in mediating the effects of cannabinoids. Activation of the endogenous cannabinoid type 1 (CB1) and type 2 (CB2) receptors has been shown to lead to the inhibition of tumor progression ^9^. CBD does not interact efficiently with CB1R and CB2R, but cannabigerol (CBG), another non-psychotropic cannabinoid that interacts with specific targets involved in carcinogenesis, is a weak partial agonist of both CB1R and CB2R ^10^. Cannabinoids also target transient receptor potential (TRP) channels ^11^, including TRPV1 ^12^ and TRPV2 ^13^, the cyclooxygenase (COX)-1 and COX-2 (also known as prostaglandin-endoperoxide synthase 2, or PTGS2) enzymes ^14^, the fatty acid amide hydrolase (FAAH) enzyme ^11^; the 5-hydroxytryptamine receptor 1A (5-HT1A) and α2-adrenergic receptor ^10^; the nuclear receptor peroxisome proliferator-activated receptor-gamma (PPAR-γ) ^15^; and the orphan G-protein-coupled receptor 55 (GPR55) ^16^.

### CB1R and CB2R

Many of the effects produced by THC and its synthetic cousins depend on their ability to target cannabinoid receptors ^17-19^, which include the GPCRs CB1R ^20^ and CB2R ^21^.

CB1Rs are located primarily in central and peripheral neurons; CB2Rs, predominantly in immune cells. CB1Rs are also expressed by some non-neuronal cells, including immune cells, and CB2Rs by some neurons both within and outside the brain (reviewed in ^22^).

CBD has very low affinity for either CB1R or CB2R ^23^. However, there is evidence that despite this low affinity for CB1R and CB2R, CBD can interact with these receptors at reasonably low concentrations ^22^, yielding such varied effects as GPR55 antagonism ^24,25^, activation of TRPV1 ^26^, and decreased cancer cell proliferation and increased neuroprotection ^18,27^.

### FAAH

FAAH is an intracellular enzyme that catalyzes the hydrolysis of the endogenous cannabinoid ligand anandamide (arachidonoylethanolamide, AEA) ^28,29^. CBD has been reported to inhibit both FAAH-mediated degradation of AEA ^30^ and AEA transporter activity ^31^. These findings suggest that some of the pharmacological actions of CBD might be due to inhibition of AEA degradation.

### GPR55

GPR55 is an orphan G protein-coupled receptor (GPCR), meaning it has no known ligand. The *GPR55* gene is expressed in several brain areas including the hippocampus, hypothalamus, frontal cortex, and cerebellum ^16^. The function of GPR55 in the periphery is varied: vasodilatation, cellular proliferation and migration, bone dynamics, energy balance, gastrointestinal processes, and inflammation and pain (reviewed in ^16^). In the central nervous system, studies suggest GPR55 could modulate procedural memory, motor coordination, anxiety, and hippocampal release of glutamate (reviewed in ^16^).

GPR55 can be activated by endocannabinoids, though the extent to which the receptor can be fully activated is still unknown ^25^. Despite poor sequence homology, many studies have reported significant functional interaction between GPR55 and CB1R/CB2R ^32-34^. As such, GPR55 is being considered as the cannabinoid receptor type 3 (CB3R) by the International Union of Basic and Clinical Pharmacology Committee on Receptor Nomenclature and Drug Classification ^32,35^. Given its role in motor coordination, modulation of GPR55 activity has been proposed as a therapeutic target for epilepsy. CBD was recently approved by the US FDA as a treatment for certain childhood forms of epilepsy (Epidiolex^®^, GW Pharmaceuticals), leading to the hypothesis that one mechanism of action of CBD is GPR55 inhibition ^25^.

### Id-1

CBD has been shown to reduce the expression of the inhibitor of differentiation and DNA binding 1 (Id-1). Id family members negatively regulate the activity of basic helix–loop–helix proteins, which are critical components of developmental processes such as sex determination and neurogenesis ^36^. Constitutive expression of Id proteins can inhibit the differentiation of various tissues ^37,38^. Perhaps most relevant, the Id proteins have been shown to be involved in the pathogenesis of human cancers ^39,40^.

The level of expression of the *ID1* gene has been shown to be strongly correlated to the invasive and metastatic behavior of breast cancer ^41,42^ and glioblastoma (GBM) ^36^ cells. Higher levels of *ID1* have been documented for a variety of aggressive tumor types including melanoma and cancers of the head and neck (squamous cell carcinoma [SCC]), esophagus (SCC), oral cavity (SCC), liver, colon, pancreas, thyroid, ovary, cervix, endometrium, and prostate (reviewed in ^39^), especially when compared to normal cells of the same tissue origin ^39,40^.

### PTGS2

The *PTGS2* gene encodes the COX-2 enzyme, which is one of two cyclooxygenases in humans. It is involved in the conversion of arachidonic acid to prostaglandin H2, an important precursor of prostacyclin, which is expressed in inflammation. PTGS2 is not expressed in most cells under normal conditions, but elevated levels are found during inflammation. Non-steroidal anti-inflammatory drugs (NSAIDs) inhibit prostaglandin production by both PTGS1 (COX-1) and PTGS2.

The expression of *PTGS2* is upregulated in many cancers. Inhibiting *PTGS2* expression may therefore be beneficial in preventing and treating certain types of cancer ^43^.

COX-2 plays a significant role in intraneuronal CB1R signal transduction and modulation of *PTGS2* and *CNR1* gene expression ^44^. CBD and Δ9-THC have been found to increase expression and activity of PTGS2, with *PTGS2* being one of the 25 most up-regulated genes upon treatment with either cannabinoid in T98G (but not U-87 MG) cells ^45^.

### TRPV1 and TRPV2

TRP channels are pore-forming transmembrane proteins that allow ions and xenobiotics to permeate biological membranes ^46^. It has been suggested that TRP channel-expressing cancer cells may allow entry of charged molecules ^47-50^ and certain anti-proliferative and cytotoxic molecules ^47^ by permeation.

CBD has been found to be a full, though weak, agonist of human TRPV1 ^26^. The TRPV2 channel is triggered by agonists such as Δ9-THC and CBD ^51,52^.

Given the importance of each of these genes to normal and pathological processes, we sought to assess the level of each across a panel of human cell lines.

## Methods

### Cell lines

The GBM lines A-172, T98G, and U-87 MG; pancreatic cancer lines MIA PaCa-2 and PANC-1; actinic keratosis line HT 297.T; melanoma lines A-375 and A101D; and normal skin fibroblasts BJ and Detroit 551 were all obtained from the American Tissue Culture Collection (ATCC) and grown according to the manufacturer’s recommendations. All assays were performed using cells that had been passaged fewer than 20 times upon arrival.

The SCC (skin) line HSC-5 ^53^ was obtained from the Japanese Collection of Research Biosources Cell Bank (JCRB) and Sekisui XenoTech.

### RNA extraction

RNA from cell samples was generated using the RNeasy Mini Kit (Qiagen) according to the manufacturer’s instructions.

### Generation of complementary DNA (cDNA)

RNA was converted to cDNA using the RT^2^ First Strand Kit (Qiagen) according to the manufacturer’s instructions.

### Quantitative PCR (qPCR)

The expression of *ACTB* (β-actin), *CNR1* (CB1R), *CNR2* (CB2R), *FAAH, GAPDH, GPR55, ID1* (Id-1), *PTGS2, TRPV1, TRPV2*, was measured by qPCR using validated human primer sets (RT^2^ qPCR Primer Assays, Qiagen) and the RT^2^ SYBR Green qPCR Mastermix (Qiagen). Values for gene expression were determined using the ΔCt method, where the cycle number threshold (Ct) of either glyceraldehyde 3-phosphate dehydrogenase (GAPDH) or β-actin (as housekeeping genes) is subtracted from the Ct value of each gene. Expression levels for the housekeeping gene transcripts were normalized to one million copies.

## Results

### Glioblastoma

We compared the level of expression of the set of cannabinoid-related genes across three different GBM lines, A-172, T98G, and U-87 MG. A-172 cells were the only cells of those assessed that had measurable expression of all genes tested. Two of the GBM lines tested, A-172 and U-87 MG, exhibited the highest expression of *ID1* among all the genes analyzed (**Error! Reference source not found**.**A** and **Error! Reference source not found**.**C**, respectively), as well as having fairly robust expression of *TRPV1*. T98G cells had the highest expression of *GPR55*, followed by *ID1* (**Error! Reference source not found**.**B**). A-172 and U-87 MG cells also exhibited robust expression of *PTGS2*, while only U-87 MG cells had robust expression of *TRPV2*.

### Pancreatic cancer

The pancreatic cancer line MIA PaCa-2 had observable and robust expression of all cannabinoid-related genes except *TRPV2* (**Error! Reference source not found**.**A**). PANC-1 cells, on the other hand, were fairly low expressers of all genes except *FAAH, ID1*, and *TRPV1* (**Error! Reference source not found**.**B**).

### Skin

The malignant melanoma line A-375 had observable expression of all cannabinoid-related genes except *TRPV2* (**Error! Reference source not found**.**A**). The genes with the highest levels of expression were *CNR1, GPR55, ID1*, and *TRPV1*. Another melanoma line, A101D, had observable expression of all genes (**Error! Reference source not found**.**B**). It, too, had *ID1* as the highest-expressing gene.

The SCC line HSC-5 (**Figure 4 Error! Reference source not found**.) and actinic keratosis line HT 297.T (**Figure 5 Error! Reference source not found**.) both had observable expression of *FAAH, ID1, PTGS2*, and *TRPV1;* HSC-5 cells also expressed *CNR1*. Similar to other lines, ID1 expression was the highest of all genes tested.

**Figure 1.**
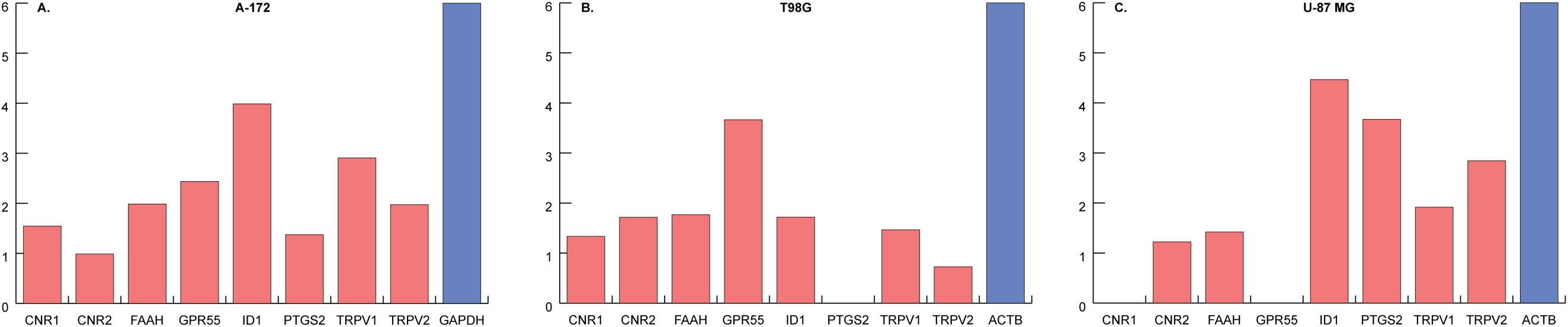
Expression of cannabinoid-related genes in glioblastoma cells. Expression levels in (**A**) A-172 cells, (**B**) T98G cells, and (**C**) U-87 MG cells were normalized to one million copies of either GAPDH (**A**) or β-actin (**B, C**).

**Figure 2.**
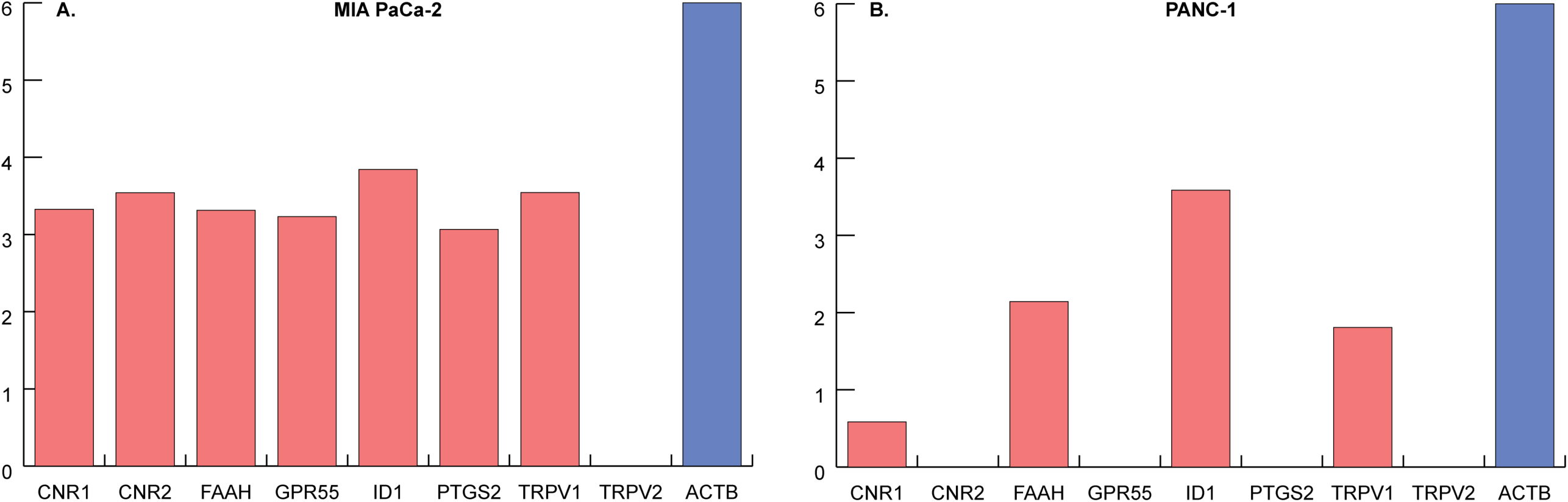
Expression of cannabinoid-related genes in pancreatic cancer cells. Expression levels in (**A**) MIA PaCa-2 cells and (**B**) PANC-1 cells were normalized to one million copies of β-actin.

**Figure 3.**
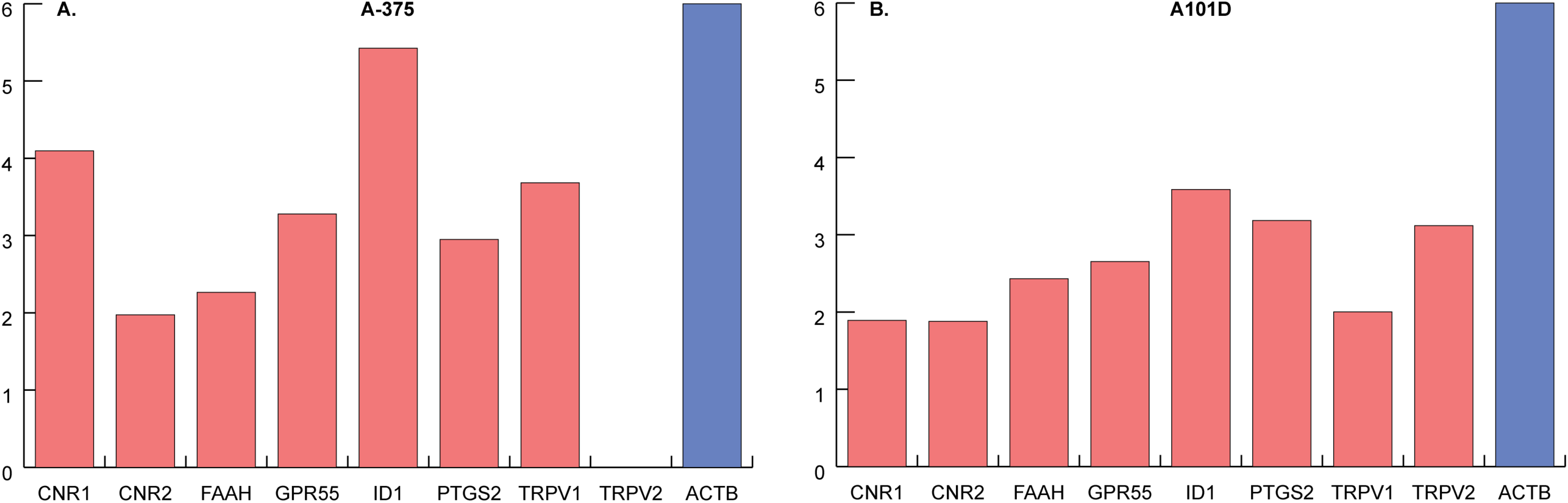
Expression of cannabinoid-related genes in melanoma cells. Expression levels in (**A**) A-375 cells and (**B**) A101D cells were normalized to one million copies of β-actin.

**Figure 4.**
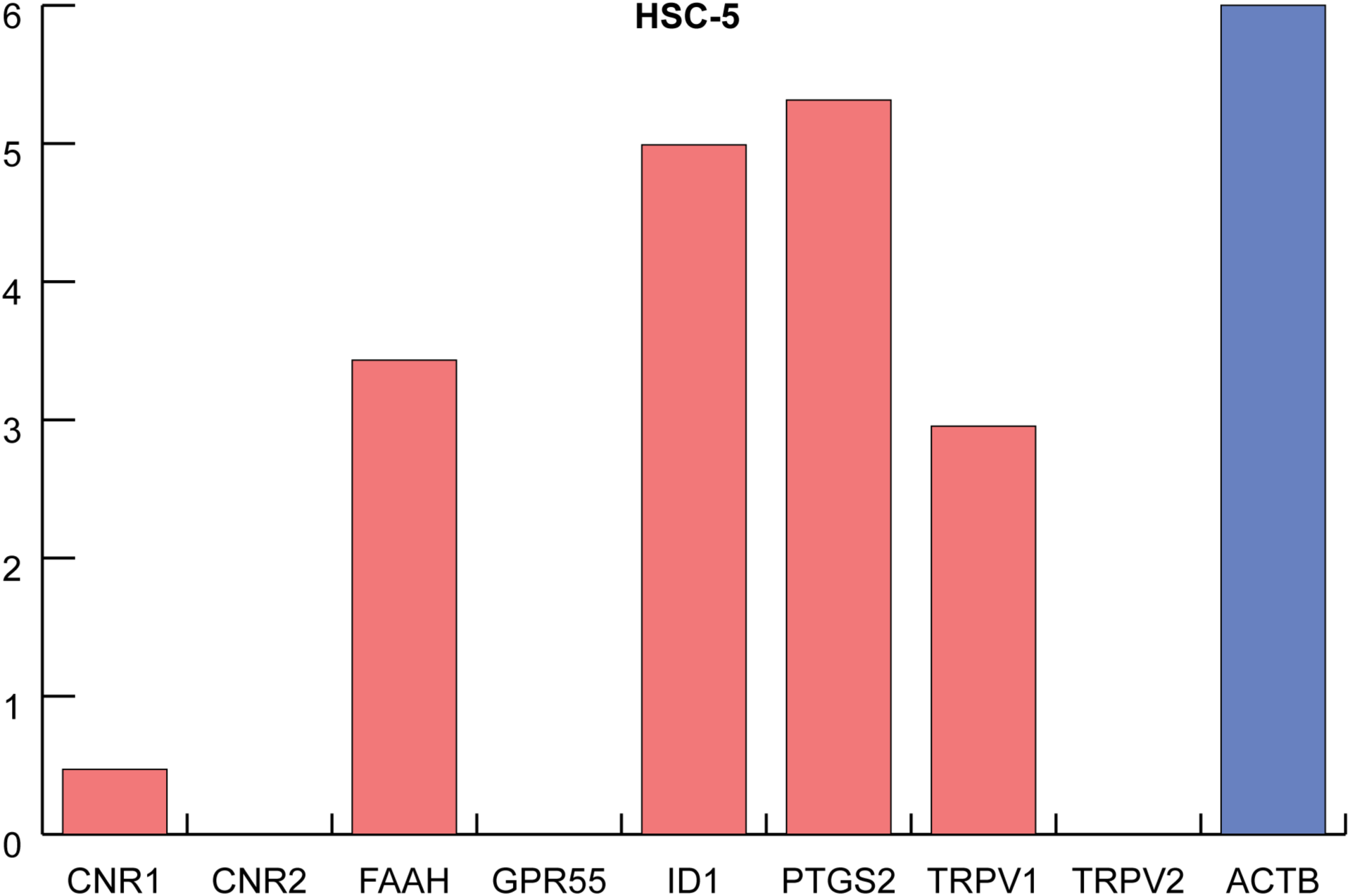
Expression of cannabinoid-related genes in HSC-5 cells, a squamous cell carcinoma line derived from skin. Expression levels were normalized to one million copies of β-actin.

**Figure 5.**
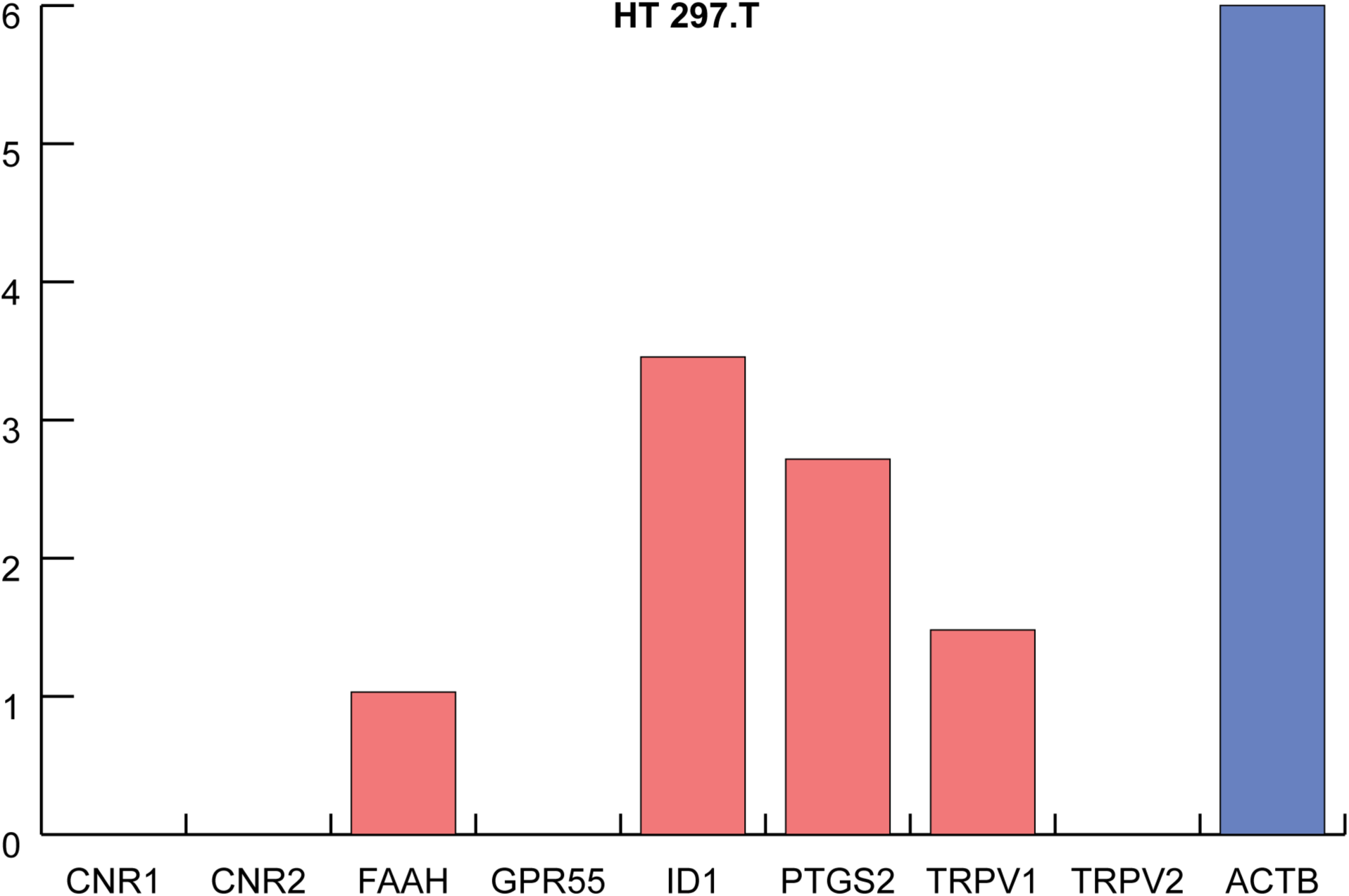
Expression of cannabinoid-related genes in HT 297.T cells, an actinic keratosis line. Expression levels were normalized to one million copies of β-actin.

We also examined gene expression in normal skin fibroblasts. The two lines studied, BJ and Detroit 551, exhibited different patterns of expression. BJ cells had high levels of *PTGS2* and *TRPV2* expression (**Error! Reference source not found**.**A**), while Detroit 551 cells had high levels of *ID1* and *PTGS2* (**Error! Reference source not found**.**B**). interestingly, BJ cells had completely undetectable levels of *ID1*.

A summary of the expression levels of the cannabinoid-related genes can be found in **Table 1**. A heatmap visualization of the same data can be found in **Error! Reference source not found**..

**Table 1.**
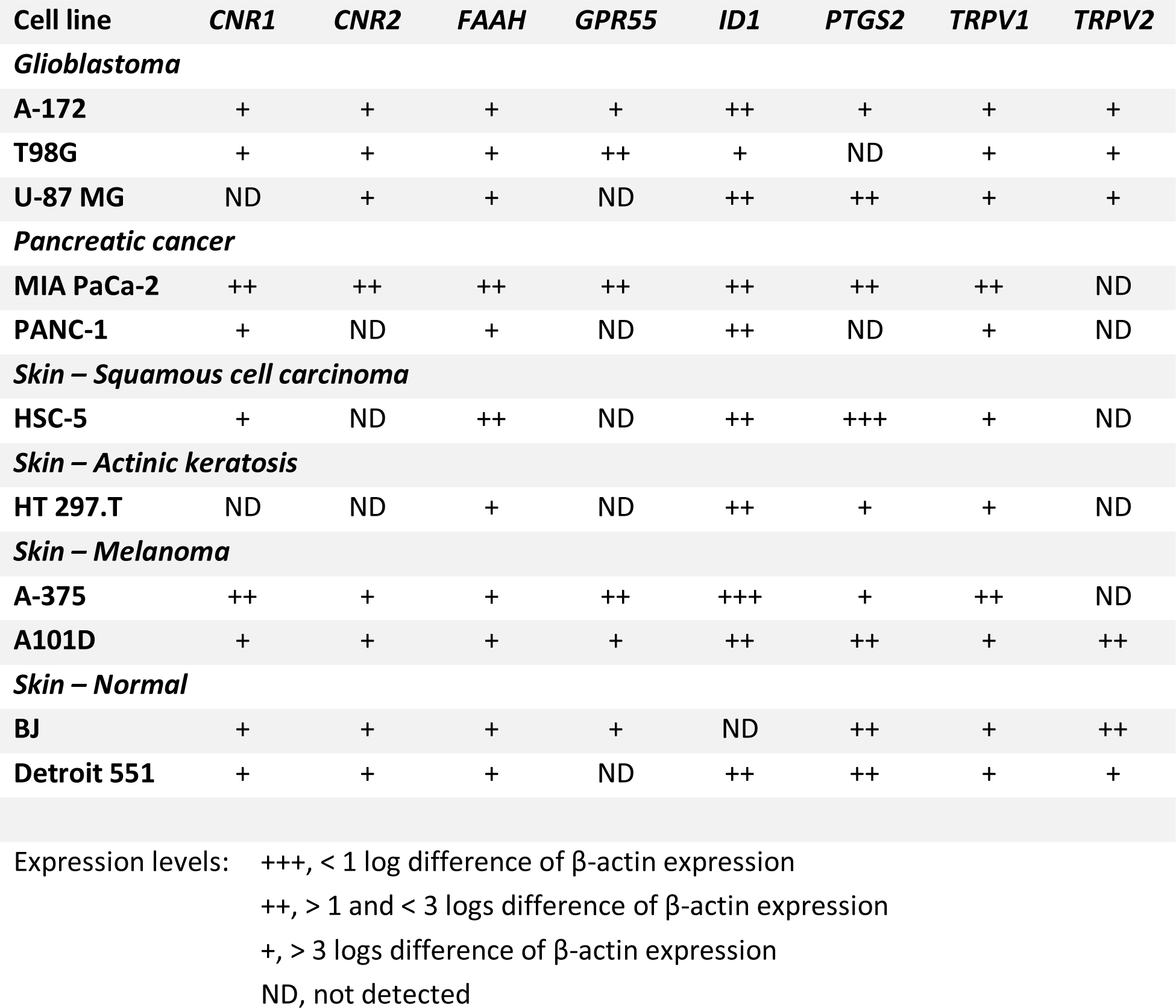
Expression of cannabinoid-related genes across multiple cell lines.

| Cell line | *CNR1* | *CNR2* | *FAAH* | *GPR55* | *ID1* | *PTGS2* | *TRPV1* | *TRPV2* |
| --- | --- | --- | --- | --- | --- | --- | --- | --- |
| ***Glioblastoma*** |  |  |  |  |  |  |  |  |
| **A-172** | + | + | + | + | ++ | + | + | + |
| **T98G** | + | + | + | ++ | + | ND | + | + |
| **U-87 MG** | ND | + | + | ND | ++ | ++ | + | + |
| ***Pancreatic cancer*** |  |  |  |  |  |  |  |  |
| **MIA PaCa-2** | ++ | ++ | ++ | ++ | ++ | ++ | ++ | ND |
| **PANC-1** | + | ND | + | ND | ++ | ND | + | ND |
| ***Skin – Squamous cell carcinoma*** |  |  |  |  |  |  |  |  |
| **HSC-5** | + | ND | ++ | ND | ++ | +++ | + | ND |
| ***Skin – Actinic keratosis*** |  |  |  |  |  |  |  |  |
| **HT 297.T** | ND | ND | + | ND | ++ | + | + | ND |
| ***Skin – Melanoma*** |  |  |  |  |  |  |  |  |
| **A-375** | ++ | + | + | ++ | +++ | + | ++ | ND |
| **A101D** | + | + | + | + | ++ | ++ | + | ++ |
| ***Skin – Normal*** |  |  |  |  |  |  |  |  |
| **BJ** | + | + | + | + | ND | ++ | + | ++ |
| **Detroit 551** | + | + | + | ND | ++ | ++ | + | + |

Expression levels: +++ , < 1 log difference of β-actin expression ++ , > 1 and < 3 logs difference of β-actin expression + , > 3 logs difference of β-actin expression ND, not detected

When visualized in the aggregate, all cell types expressed cannabinoid-related genes, though to varying degrees (**Figure 8 Error! Reference source not found**.). The highest total gene-expressing cell line was MIA PaCa-2, while the highest gene-expressing cell type was melanoma.

**Figure 6.**
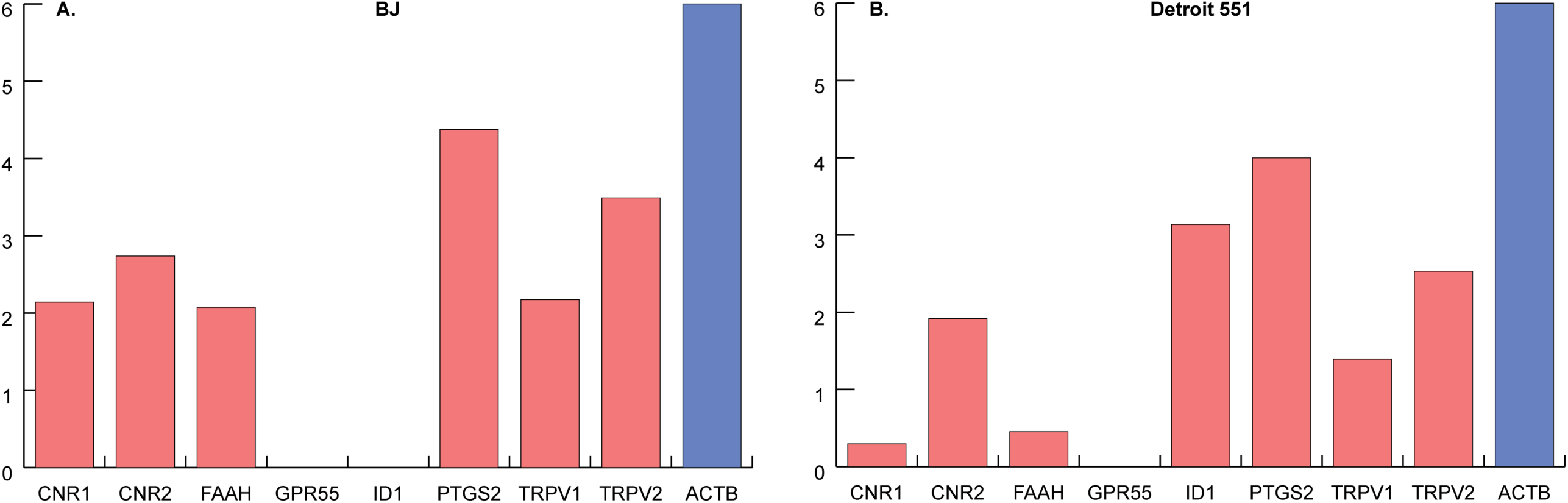
Expression of cannabinoid-related genes in normal skin fibroblasts. Expression levels in (**A**) BJ cells and (**B**) Detroit 551 cells were normalized to one million copies of β-actin.

**Figure 7.**
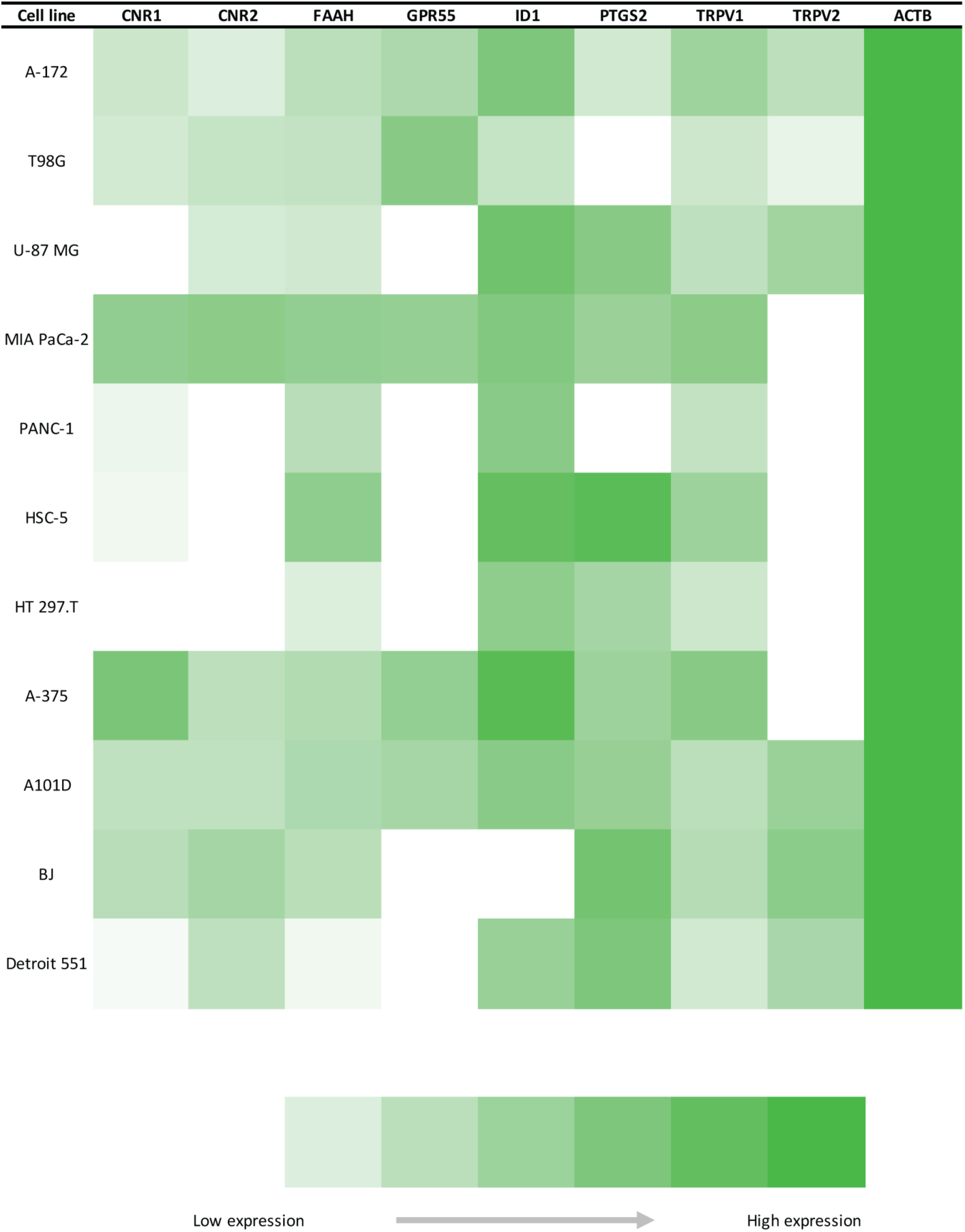
Heatmap depicting the expression of cannabinoid-related genes across multiple cell lines. The colors indicate levels of gene expression and range from white (nadir of expression) to dark green (zenith of expression).

**Figure 8.**
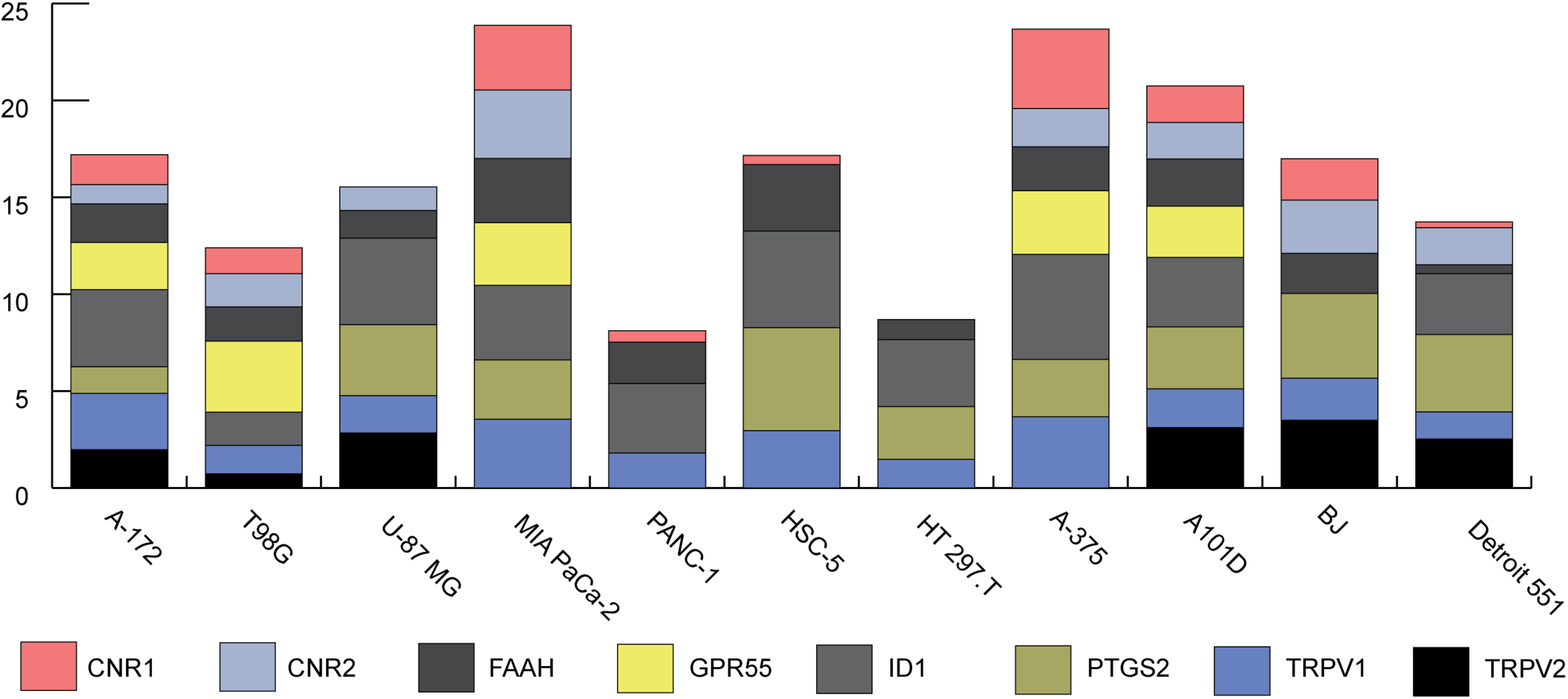
Expression of cannabinoid-related genes in normal skin fibroblasts. Expression levels in (**A**) BJ cells and (**B**) Detroit 551 cells were normalized to one million copies of β-actin.

### ID1 expression

Because of the special role Id-1 appears to play in tumorigenesis, and the fact that CBD has been shown to down-regulate *ID1* gene expression and associated glioma cell invasiveness and self-renewal ^36^, we paid particular attention to its expression levels in the cell lines assessed in this study. We found that expression of *ID1* was consistently high in cancerous cell lines (**Figure 9**Error! Reference source not found.). The only other diseased line available, actinic keratosis, had the lowest levels of *ID1* expression. Two normal skin fibroblast lines, BJ and Detroit 551, showed different levels of *ID1* expression, with BJ having undetectable *ID1* levels.

**Figure 9.**
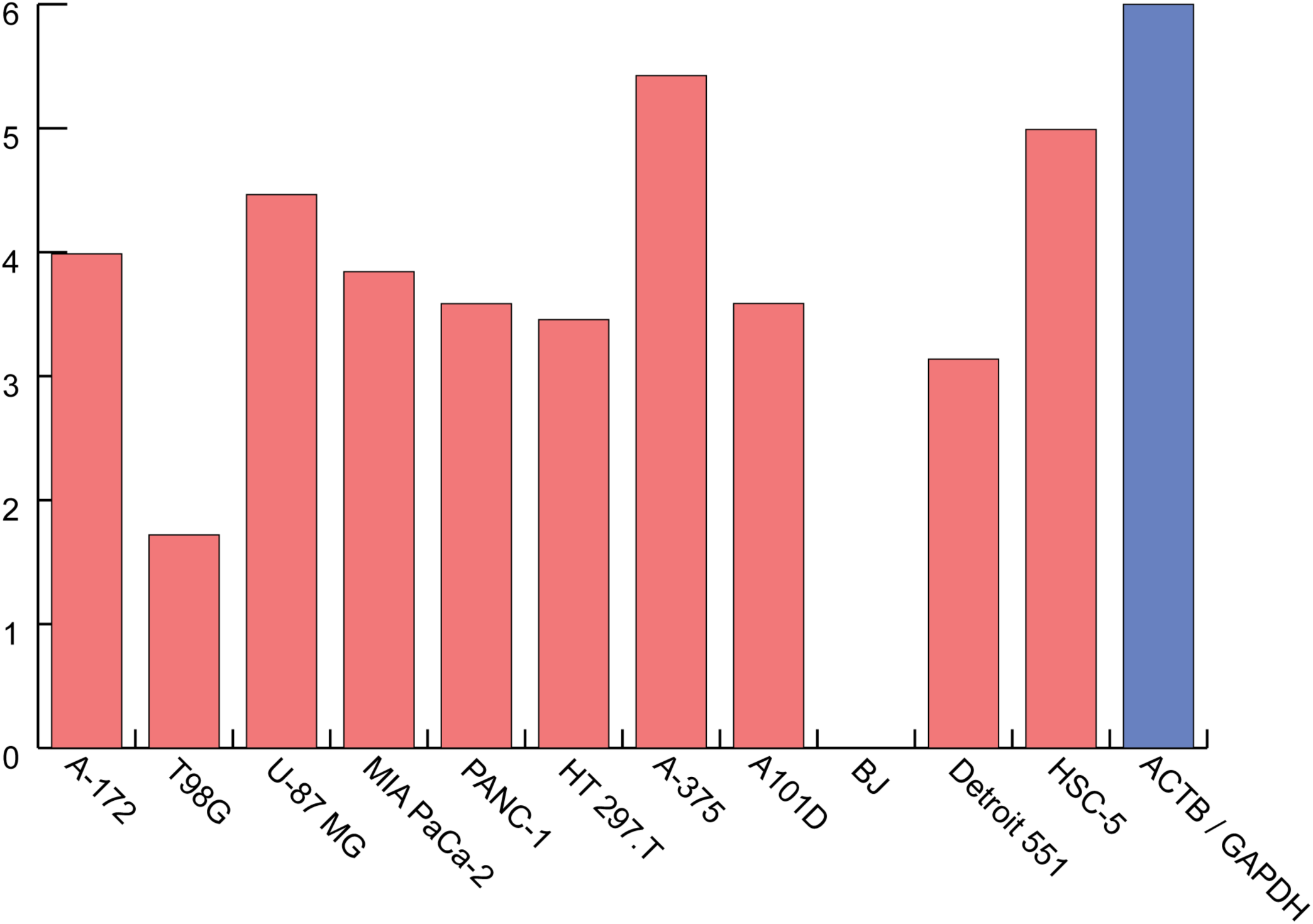
ID1 mRNA expression in different cell lines. Expression levels were normalized to one million copies of the housekeeping gene (β-actin or GAPDH).

## Discussion

Cannabinoids, and CBD in particular, have been proposed as therapeutic agents for a wide variety of diseases. Although the precise mechanism(s) of action by which cannabinoids exert their effects remain(s) unknown, it has been suggested that the endogenous expression of enzymes, receptors, signaling molecules, and other mediators that comprise the endocannabinoid system render the human body capable of functional responses to this class of compounds. To our knowledge, this study is the first systematic assessment of cannabinoid-related genes across multiple cell lines that span human diseases.

Virtually all of the cell lines we tested exhibited robust expression of *ID1*, a gene that has been implicated in the pathogenesis of human cancers ^39,40^. (U-87 MG cells were previously shown to be negative for *ID1* expression ^54^.) As CBD has been shown to reduce the expression of *ID1*, thereby reducing either the likelihood of cellular transformation or the aggressiveness of existing tumors, these results suggest therapeutic targeting of these diseases with CBD may improve outcomes.

The expression of *PTGS2* was second only to *ID1* across the cell lines tested. Given its role in mediating inflammation, it was not surprising to observe fairly high levels in transformed cell lines (even if some are characterized as “normal”). This observation is consistent with the finding that *PTGS2* expression is upregulated in many cancers. What is counter-intuitive, however, is the observation that CBD was found to up-regulate expression of *PTGS2*, as well as increase the activity of the COX-2 enzyme ^45^. Increasing the activity of a pro-inflammatory mediator such as COX-2 contradicts the narrative that CBD and other cannabinoids are potent anti-inflammatory drugs, exerting their effects through induction of apoptosis, inhibition of cell proliferation, suppression of cytokine production, and induction of regulatory T cells ^5^. Further investigation into the effect cannabinoid treatment of these cell lines has on *PTGS2* expression is warranted.

CBD has very low affinity for either CB1R or CB2R ^23^. In addition, CBD has been reported to inhibit both FAAH-mediated degradation of the endogenous cannabinoid ligand, anandamide ^30^ and the RBL-2H3 cell anandamide transporter activity ^31^. These findings raised the possibility that some of CBD’s pharmacological actions might be due to inhibiting the degradation of anandamide. Anandamide has been identified as an endogenous CB1R agonist and as a ligand for CB2R (despite low affinity).

A comprehensive survey of gene expression in human tissues was conducted by RNA sequencing (RNA-seq) ^55^. Among 27 different human tissues assessed by RNA-seq, *CB1R* expression was highest in the brain; skin was fairly low, pancreas was virtually undetectable. *CB2R* expression, highest in the lymph nodes, was virtually undetectable in the brain, pancreas, and skin. *FAAH* expression, highest in the prostate, was approximately equivalent in the brain and skin and twice that in the pancreas. *GPR55* expression was highest in the testis; expression in the brain and skin was approximately the same, with virtually undetectable levels in the pancreas. *ID1* expression was highest in the colon and lung, with intermediate expression in the brain and skin, and virtually undetectable levels in the pancreas. *PTGS2* expression, highest in the bone marrow, was low in the brain and virtually undetectable in the pancreas and skin. *TRPV1* expression was highest in the small intestine; expression in the skin was approximately twice that in the brain and fourfold that in the pancreas. Finally, *TRPV2* expression was highest in the lung; expression in the brain and skin was approximately the same, and fivefold that in the pancreas.

In cancer, the situation is somewhat different. *CB1R* and *CB2R* are both overexpressed in several tumors, including pancreatic adenocarcinoma ^56^, suggesting that a cannabinoid-based therapy may activate cell death pathways predominantly in tumor cells. In cell lines, our results are somewhat different than those reported previously ^57^. In the GBM line T98G, the authors found low expression of *GPR55, TRPV1*, and *TRPV2*, and undetectable levels of *CB1R* and *CB2R*; we report T98G cells as expressing intermediate levels of *GPR55*, and low levels of the other three genes. For another GBM line, U-87 MG, found low levels of all genes except *CB2R*, which was undetectable; similarly, we report intermediate to low levels of each gene. In the malignant melanoma line A-375, Baram *et al*. reported high levels of *CB1R* and *<u>TRPV2</u>*; in our study, we confirm fairly high levels of *CB1R* but did not detect *TRPV2* expression.

Another important site at which anandamide acts is TRPV1 ^55^, which acts as a molecular integrator of noxious stimuli and is important for thermal nociception and inflammatory hyperalgesia ^56,57^. CBD has been shown to be a full, although weak, agonist of human TRPV1 ^26^. Thus, CBD could activate either cannabinoid receptors indirectly through anandamide or vanilloid receptors directly or indirectly via anandamide.

In conclusion, the cell types assessed in this study – diseased and normal cell types, cancer and non-cancerous cell types – all exhibited expression of one or more gene products that have been shown to mediate the pharmacological effects of cannabinoids. Moreover, the diversity of these cell types is consistent with the notion that our bodies, whether healthy or sick, are innately capable of responding to cannabinoids. Although this paper reports levels of RNA transcripts, we cannot conclude that the expression patterns are similar for the corresponding proteins; further investigation is therefore warranted. Our findings strongly suggest, however, there is a potential therapeutic role for phytocannabinoids in the context of disease.

## Acknowledgements

The authors wish to thank Sophia Podber, Everest Maya-Tudor, and Soyeon Lee for their important contributions to this study.

